# N-terminal tags impair the ability of Lamin A to provide structural support to the nucleus

**DOI:** 10.1101/2024.04.19.590311

**Authors:** Jacob Odell, Jan Lammerding

**Affiliations:** Weill Institute for Cell and Molecular Biology, Cornell University, Ithaca, NY 14853; Graduate Field of Biochemistry, Molecular and Cell Biology, Cornell University, Ithaca, NY 14853; Meinig School of Biomedical Engineering, Cornell University, Ithaca, NY 14853

**Keywords:** nucleus, lamins, fluorescent tags, emerin, GFP, mechanobiology

## Abstract

Lamins are intermediate filament proteins that contribute to numerous cellular functions, including nuclear morphology and mechanical stability. The N-terminal head domain of lamin is critical for higher order filament assembly and function, yet the effects of commonly used N-terminal tags on lamin function remain largely unexplored. Here, we systematically studied the effect of two differently sized tags on Lamin A (LaA) function in a mammalian cell model engineered to allow for precise control of expression of tagged lamin proteins. Untagged, FLAG-tagged, and GFP-tagged LaA completely rescued nuclear shape defects when expressed at similar levels in lamin A/C-deficient (*Lmna^−/−^*) MEFs, and all LaA constructs prevented increased nuclear envelope (NE) ruptures in these cells. N-terminal tags, however, altered the nuclear localization of LaA and impaired the ability of LaA to restore nuclear deformability and to recruit Emerin to the nuclear membrane in *Lmna^−/−^* MEFs. Our finding that tags impede some LaA functions but not others may explain the partial loss of function phenotypes when tagged lamins are expressed in model organisms and should caution researchers using tagged lamins to study the nucleus.

## Introduction

Lamins are nuclear intermediate filament proteins that form a dense protein meshwork underlying the inner nuclear membrane (Aebi et al., 1986; Buxboim et al., 2023; Shimi et al., 2015). Lamins have many important functions in metazoan cells, where they provide structural support to the nucleus, regulate nuclear morphology, and influence gene expression (de Leeuw et al., 2018; Kalukula et al., 2022; Vahabikashi et al., 2022). Research on lamins frequently involves the fusion of a genetically encoded fluorophore or epitope tag to the lamin protein to allow for detection in live cells and other assays. Lamins are sensitive to where the tag is placed within the protein sequence (Herrmann et al., 2002); C-terminal tags lead to obvious loss of function phenotypes, as the tag disrupts the lamin C-terminal CaaX motif important for proper post-translational processing and localization of lamins (Kreplak et al., 2008), and the end of prelamin A is cleaved during processing into mature Lamin A (hereafter referred to as LaA) (Sinensky et al., 1994). Consequently, the vast majority of studies use N-terminal tags to label LaA. For example, N-terminally tagged lamins have been used to visualize the localization and dynamics of lamins in live cells throughout the cell cycle (Broers et al., 1999), study interactors of lamins (Roux et al., 2012), investigate autophagy (Dou et al., 2015), and serve as a marker of the nucleus in live cells and organisms (Morin et al., 2001; Rizzo et al., 2009).

In vivo studies using tagged lamins suggest that the presence of a tag can be detrimental for proper lamin assembly and function. In the nematode *C. elegans*, endogenous tagging of its lone lamin with GFP at the N-terminus causes mild or moderate fitness defects (Bone et al., 2016; Gregory et al., 2023) and abnormal lamina assembly and stability (Velez-Aguilera et al., 2020). GFP-tagged lamins have also been associated with impaired function in *Drosophila*; endogenously tagged GFP-LamC marks the NE as expected, but leads to bright nuclear granules (Morin et al., 2001; Schulze et al., 2005) and abnormal chromatin patterns (Gurudatta et al., 2010), suggesting that the chimeric protein has compromised function. However, no systematic comparison of differently sized N-terminal tags on mammalian lamins has been performed to date, and many of the early studies using fluorescently tagged mammalian lamins involved overexpression on top of endogenous lamins, which may mask loss-of-function effects of the tag.

To address these knowledge gaps, we determined the effects of differently sized N-terminal tags on LaA function in a mammalian cell model, comparing the ability of untagged and tagged LaA constructs to rescue various LaA functions in lamin A/C-deficient (*Lmna^−/−^*) mouse embryo fibroblasts (MEFs). Specifically, we examined differences in the rescue potential of LaA with a small N-terminal tag (FLAG: 8 amino acids, 1 kDa) or large N-terminal tag (enhanced GFP: 238 amino acids, 27 kDa) compared to untagged LaA and assessed these proteins’ ability to rescue nuclear morphology, deformability, rupture, and disturbed localization of lamin-interacting proteins in *Lmna^−/−^*MEFs. Although tagged LaA restored some LaA functions, other functions were moderately or severely impaired by the addition of even a small tag, leading us to caution researchers about the use of tagged LaA proteins.

## Results and Discussion

*Lmna^−/−^* MEFs are one of the best characterized systems used to study the function of LaA (Chen et al., 2021, 2018; Coffinier et al., 2010; Lammerding et al., 2006, 2004; Odell et al., 2024; Sullivan et al., 1999; Zwerger et al., 2013), and nuclear defects can be rescued by re-introduction of human LaA (Lammerding et al., 2006; Odell et al., 2024; Wintner et al., 2020). Thus, we genetically modified *Lmna^−/−^* MEFs to allow for doxycycline inducible expression of untagged human LaA, FLAG [DYKDDDDK]-LaA, or GFP-LaA and then derived clonal populations with tunable LaA expression levels (Figure 1A-B). Precisely controlling the expression levels of the exogenously expressed lamin constructs is crucial for quantitative comparison, since previous studies have shown that LaA expression levels directly correlate with nuclear stiffness (Lammerding et al., 2006; Srivastava et al., 2021; Swift et al., 2013) and that severe overexpression of GFP-tagged lamins can cause aggregation and mislocalization (Rizzo et al., 2009). Note that we did not add any additional linker residues to the tags, i.e., FLAG or GFP sequences were added directly adjacent to the LaA sequence. We identified individual clones and specific doxycycline concentrations that yielded similar levels of LaA expression across the differently tagged constructs and that resulted in expression levels comparable to wild-type LaA in *Lmna*^+/+^ MEFs (Figure 1B-D).

**Figure 1:**
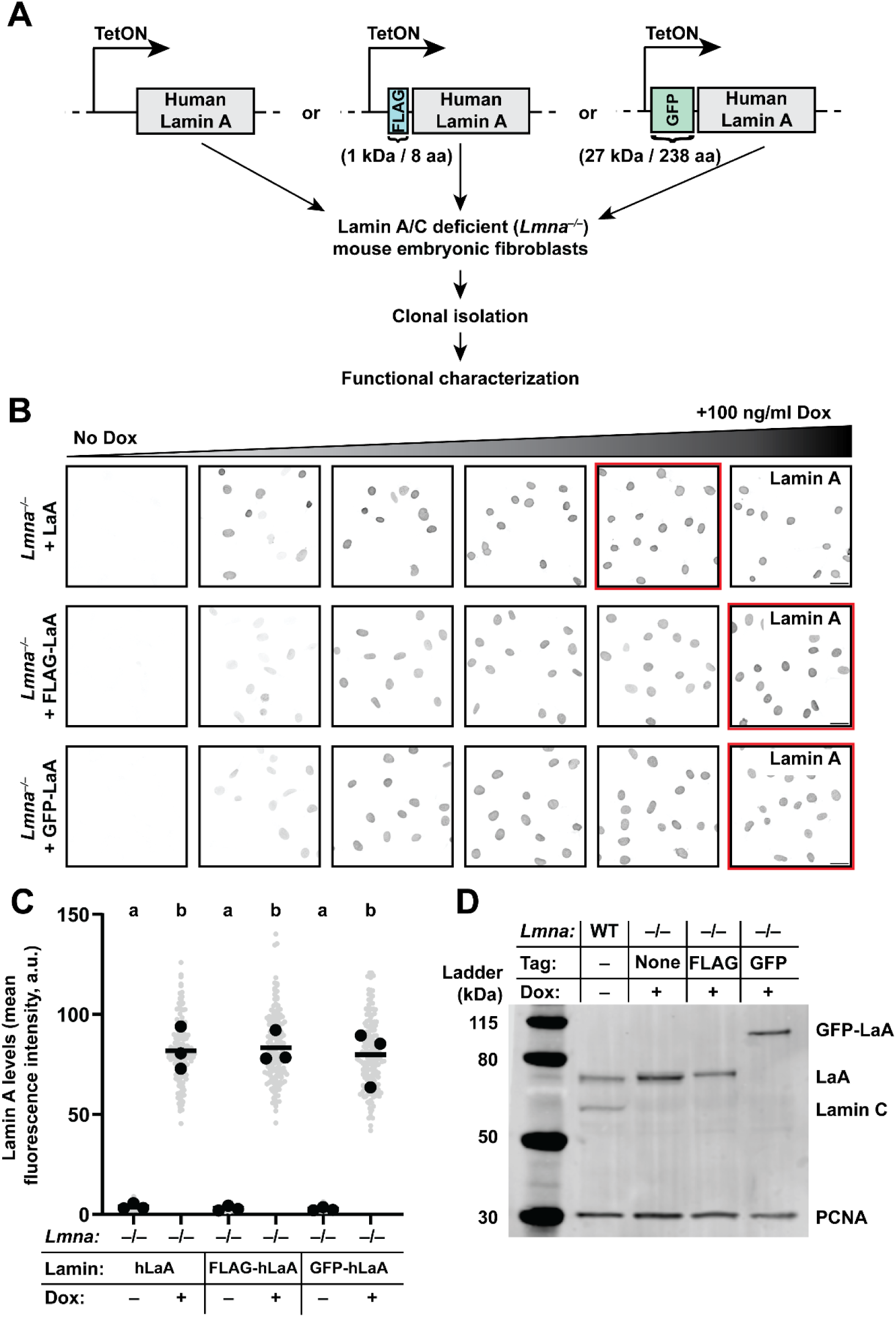
Expressing tagged LaA constructs at defined levels. (**A**) Overview of lamin constructs, exogenous expression system, and experimental pipeline to compare the effect of different N-terminal tags on LaA function. (**B**) LaA expression in clonal *Lmna^−/−^*MEFs following titration of doxycycline levels. Conditions that resulted in similar levels of LaA expression are highlighted by a red bounding box. Scale bar: 50 µm. (**C**) Quantification of nuclear LaA levels by immunofluorescence staining following treatment with concentrations of doxycycline identified in (**B**). One-way ANOVA was performed on replicate means (black points), and Tukey’s multiple comparison test was used to compare each group to every other group. Sets of points with the same letter above them are not significantly different, whereas different letters indicate *p* < 0.05. (**D**) Immunoblot analysis showing similar levels of LaA expression between the different cell lines/constructs, following treatment with doxycycline levels identified in (**B**). PCNA was used as a loading control.

### Nuclear morphology and LaA distribution in cells expressing differently tagged LaA constructs

All three LaA constructs localized to the nucleus and were enriched at the nuclear periphery as expected (Figure 2A). We previously showed that loss of lamin A/C results in abnormally shaped nuclei with decreased nuclear circularity (Lammerding et al., 2006; Odell et al., 2024; Zwerger et al., 2013). Expression of all three LaA constructs increased nuclear circularity in *Lmna^−/−^* MEFs (Figure 2B), indicating functional rescue. We did not detect any difference in the circularity of nuclei prior to inducing LaA expression, and all LaA constructs, regardless of the presence of a tag, showed similar efficacy in restoring nuclear shape to levels of wild-type cells (Figure 2B).

**Figure 2:**
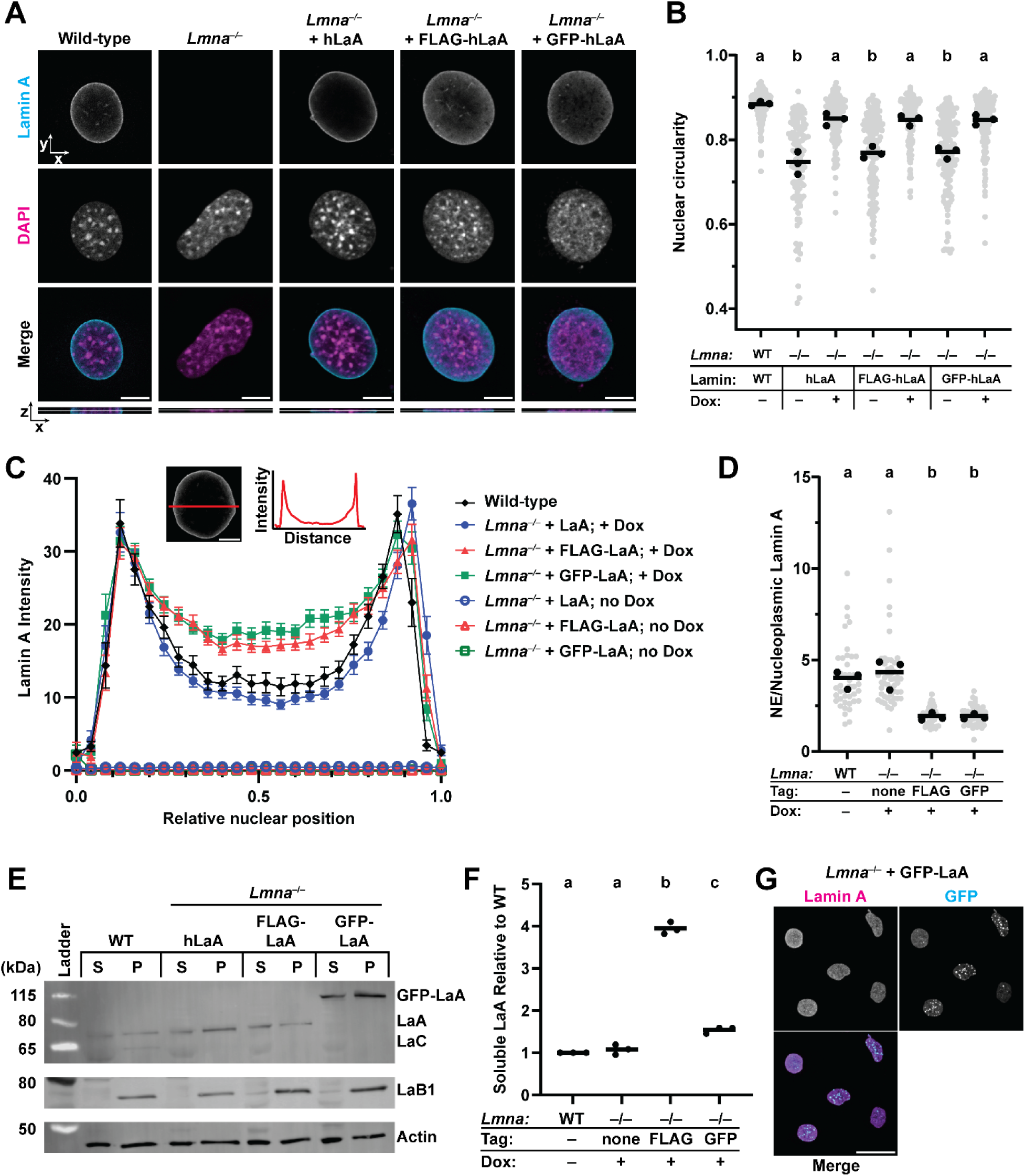
N-terminal tags interfere with proper localization of LaA. (**A**) Representative images of cells immunofluorescently labeled for LaA. Single confocal z-slices are shown, with the orthoview shown below each set of images. The horizontal white line through each orthoview indicates the plane shown in the main figure. Scale bar: 10 µm. (**B**) Nuclear circularity analysis of cells expressing different LaA constructs, using the doxycycline conditions identified in Figure 1. (**C**) Quantification of LaA intensity profiles along a line through the midplane of the nucleus (see red line in inset). (**D**) Quantification of LaA fluorescence intensity ratio between the NE and nucleoplasmic LaA signal for the cell lines shown in (**C**). (**E**) Solubility of LaA constructs using buffers of different stringency. S: supernatant (soluble fraction), P: pellet (insoluble fraction). (**F**) Quantification of soluble LaA relative to WT. (**G**) Representative image showing GFP puncta in some *Lmna^−/−^* MEFs expressing GFP-LaA. Scale bar: 50 µm.

Impaired lamin filament assembly can result in altered intranuclear distribution, such as loss of LaA from the NE and an increase in nucleoplasmic LaA (Manju et al., 2006; Wallace et al., 2023; Zwerger et al., 2015, 2013). To assess the effect of the different tags on intranuclear localization of LaA, we measured the intensity profiles of LaA across the nucleus following indirect immunofluorescence staining with an antibody recognizing an epitope present in all three LaA constructs (Figure 2C). The three LaA constructs were enriched at the nuclear periphery to similar extents, but both FLAG-LaA and GFP-LaA had increased nucleoplasmic localization compared to untagged LaA (Figure 2A, C, D). Furthermore, both FLAG-LaA and GFP-LaA were significantly more soluble than untagged LaA, based on extraction with different stringency buffers (Figure 2E-F). GFP-LaA also formed puncta in some nuclei, which may represent aggregations that resisted extraction by low stringency buffers (Figure 2G). These findings suggest that the addition of an N-terminal tag interferes with proper LaA assembly into the nuclear lamina, and are in agreement with previous studies, which reported defects in the assembly and/or localization when using GFP-tagged lamins (Morin et al., 2001; Velez-Aguilera et al., 2020).

### N-terminal tags interfere with LaA’s ability to rescue nuclear stiffness

Lamins are an important contributor to nuclear deformability, and loss of lamin A/C leads to nuclei with decreased nuclear stiffness (Davidson et al., 2019; Earle et al., 2020; Lammerding et al., 2006; Pajerowski et al., 2007; Wintner et al., 2020). To assess the effect of different tags on LaA’s ability to rescue nuclear stiffness in *Lmna*^−/−^ MEFs, we applied a high-throughput micropipette aspiration assay (Davidson et al., 2019) to these cells, which imposes large deformations on the nucleus (Figure 3A-B). Whereas small nuclear deformations are predominantly resisted by chromatin, large nuclear deformations are primarily resisted by lamins (Stephens et al., 2017), making this assay ideally suited to measure the mechanical function of lamins. Without induction of exogenous LaA, all *Lmna^−/−^* clonal MEF lines had similar levels of nuclear deformability, and all *Lmna^−/−^*MEF lines had more deformable nuclei than wild-type cells (Figure 3A-D). All three LaA constructs were able to significantly reduce nuclear deformability compared to the non-induced controls (Figure 3A-D). However, only untagged LaA was able to restore nuclear deformability to wild-type levels (Figure 3D). Despite their difference in size, both the FLAG-tag and the GFP-tag resulted in similar levels of (partial) rescue. These results suggest that the addition of an N-terminal tag, even as small as a FLAG-tag, inhibits the ability of LaA to regulate nuclear stiffness.

**Figure 3:**
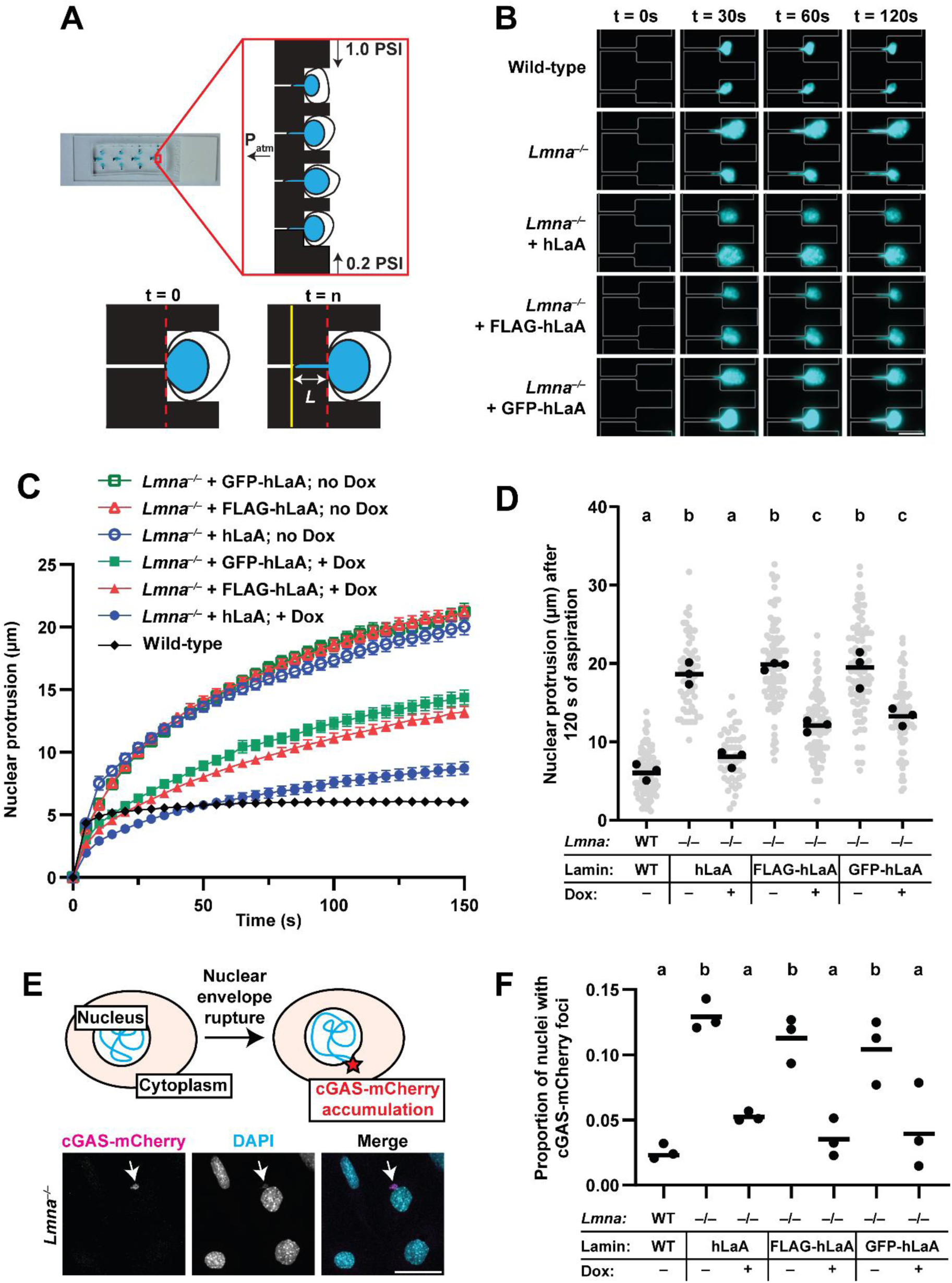
N-terminal tag impairs ability of LaA to rescue nuclear stiffness but not NE rupture. (**A**) Microfluidic micropipette aspiration assay, adapted from (Odell et al., 2024). A cell suspension is perfused into the device under a precisely controlled pressure gradient, allowing single cells to become trapped in the aspiration pockets (top).

In addition to having more deformable nuclei, cells lacking lamin A/C have increased rates of NE rupture (Cho et al., 2019; De Vos et al., 2011; Denais et al., 2016; Earle et al., 2020; Raab et al., 2016). We examined the ability of the different LaA constructs to reduce the rate of spontaneous NE rupture in *Lmna^−/−^* MEFs using a fluorescent reporter for NE rupture consisting of cyclic GMP-AMP synthase (cGAS) fused with mCherry that accumulates at sites where nuclear DNA is exposed to the cytoplasm (Figure 3E) (Denais et al., 2016; Raab et al., 2016). Consistent with previous studies (Denais et al., 2016; Odell et al., 2024; Raab et al., 2016), we found that *Lmna*^−/−^ MEFs had increased rates of spontaneous NE rupture compared to wild-type controls (Figure 3F). Expression of either untagged LaA, FLAG-LaA, or GFP-LaA reduced the NE rupture rates in *Lmna*^−/−^ MEFs (Figure 3F). Surprisingly, despite their different effects on nuclear deformability, all LaA constructs were able to restore NE rupture rates in *Lmna*^−/−^ MEFs to levels of wild-type MEFs, suggesting that the addition of an N-terminal tag does not interfere with LaA’s ability to maintain nuclear membrane integrity.

Individual cells are gradually aspirated through small (3 × 5 µm^2^ cross-section) micropipette channels along a larger pressure gradient (bottom). The rate and extent (‘*L*’) of the nuclear protrusion into the aspiration channel provides information on the mechanical properties of the nucleus. (**B**) Time-dependent nuclear protrusion during micropipette aspiration. Grey outlines indicate boundaries of micropipette channels. Scale bar: 20 µm. (**C**) Quantification of nuclear protrusion over time. Plots depict mean nuclear protrusion length from three independent experiments; error bars represent s.e.m. (**D**) Nuclear protrusion lengths at 120 s after the start of aspiration. (**E**) Schematic overview of cGAS-mCherry reporter function and corresponding representative images of *Lmna*^−/−^ MEFs expressing cGAS-mCherry reporter. Arrowhead indicates site of nuclear rupture, where a cGAS-mCherry puncta overlaps with DNA (DAPI) spilling into the cytoplasm. Scale bar: 50 µm. (**F**) Quantification of NE rupture rate, i.e., the fraction of cells with mCherry signal adjacent to a DAPI stained nucleus.

### N-terminal tags interfere with ability of LaA to localize and interact with Emerin

Emerin is a NE protein and a known LaA interactor that is retained at the inner nuclear membrane (INM) by lamin A/C (Fernandez et al., 2022; Liddane and Holaska, 2021). In *Lmna^−/−^* MEFs, Emerin was mislocalized from the NE and primarily distributed to the cytoplasm (Figure 4A-B), consistent with previous reports (Guo et al., 2014; Sullivan et al., 1999; Vaughan et al., 2001). In the absence of exogenous LaA expression, all clonal *Lmna^−/−^*MEFs had similar levels of Emerin mislocalization. Upon expression, all three LaA constructs increased the presence of Emerin at the NE, but only the untagged LaA and the FLAG-LaA constructs fully restored normal localization of Emerin (Figure 4B-C). These results suggest that the presence of the large N-terminal GFP tag interferes with proper association of LaA with Emerin, whereas the smaller FLAG tag does not disturb this interaction. To determine if the N-terminal GFP-tag impaired *in vitro* binding between LaA and Emerin, we performed co-immunoprecipitation analysis using lysates from each of the cell lines expressing different LaA constructs. Emerin co-immunoprecipitated (co-IP’ed) with untagged LaA at much higher levels than with the FLAG-LaA or GFP-LaA constructs (Figure 4D-E). In contrast, co-IPs performed using a non-specific isotype matched control IgG did not yield any of the tagged lamin constructs nor Emerin (Suppl. Figure S1). Despite FLAG-LaA being able to fully rescue nuclear Emerin localization in cells, FLAG-LaA had a limiting effect on the interaction of LaA with Emerin measured by co-IP. Notably, these orthogonal approaches probe for different abilities of LaA to bind Emerin. In cells, Emerin can interact with fully assembled LaA filaments near the nuclear membranes, whereas in cell lysates, the NE is disrupted and the lamina is broken down by detergents and reducing agents. Thus, one explanation for the differing results obtained between the immunofluorescence and co-IP experiments is that both the FLAG and GFP tags reduce LaA-Emerin binding (as shown by the co-IP), but that the smaller FLAG tag does not inhibit the interaction between Emerin and assembled filaments as severely as the GFP tag, thereby leading to improved nuclear localization of Emerin in cells expressing FLAG-LaA compared to GFP-LaA. Nonetheless, our results suggest that the presence of the bulky N-terminal GFP tag interferes with the ability of LaA to interact with Emerin when LaA is both assembled into the lamina and when the lamina is broken down.

**Figure 4:**
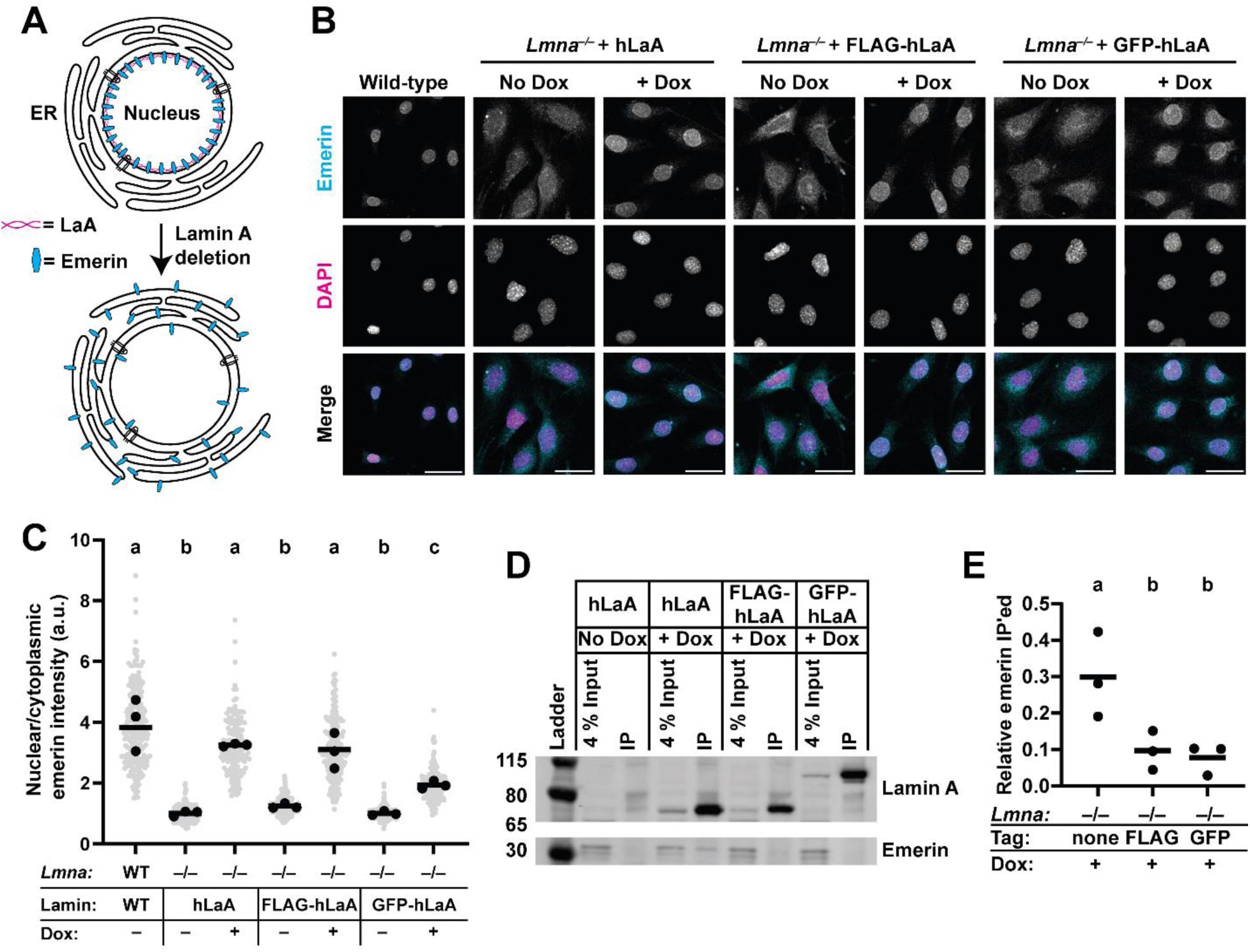
N-terminal tags interfere with the interaction of LaA with Emerin. (**A**) Emerin is mislocalized from the INM to the ER upon loss of lamin A/C. (**B**) Representative immunofluorescence images of wild type and *Lmna^−/−^* MEFs expressing different LaA constructs and stained for Emerin and DNA (DAPI). Scale bar: 50 µm. (**C**) Quantification of the mean nuclear/cytoplasmic ratios of Emerin immunofluorescence signal in individual cells. (**D**) Immunoblot of lysates from the indicated cells following IP with an anti-LaA antibody. 4% of the cell lysate was withheld from the IP experiment and ran on the gel to demonstrate equal amounts of protein in the lysates prior to IP. (**E**) Quantification of Emerin that co-IP’ed with differently tagged lamin constructs.

Taken together, our studies indicate that N-terminal tags perturb some, but not other functions of LaA. FLAG- and GFP-tagged LaA were able to completely overcome loss of endogenous lamin A/C in restoring nuclear circularity and NE rupture assays. However, N-terminal tags impaired LaA localization, rescue of nuclear deformability, and LaA interactions with Emerin. We previously showed that in iPSC-derived cardiomyocytes harboring laminopathic LaA mutations, mutant LaA was enriched in the nucleoplasm and led to reduced nuclear stiffness (Wallace et al., 2023), similar to the effects we describe here with FLAG-LaA or GFP-LaA. We propose that both mutant LaA proteins and tagged LaA constructs can alter proper assembly of lamins into higher order filaments. Structural studies have highlighted the importance of the head domain of lamin in the lateral assembly of lamin monomers into dimers, and the ultimate association into higher order filaments (Heitlinger et al., 1992; Zhou et al., 2021). By adding the highly charged FLAG tag or the globular GFP tag to the N-terminus of LaA, these motifs may interrupt the normal assembly dynamics of LaA, leading to an increased pool of unincorporated LaA. This in turn could result in a reduction in the number of filaments or altered filament and lamin network mechanics, leading to the defects observed in response to the nuclear deformability assay. Nonetheless, the networks formed by tagged lamins are sufficient to prevent NE rupture, which often occurs at sites where Lamin B1 is locally depleted or that have defects in the local nuclear lamina organization (Denais et al., 2016; Deviri et al., 2017; Pfeifer et al., 2022). We previously showed that tagged-LaA can rescue the disturbed distribution of Lamin B1 in *Lmna^−/−^*MEFs (Odell et al., 2024), supporting a mechanism where the expression of LaA in *Lmna^−/−^* MEFs improves the Lamin B network, thereby reducing NE rupture.

The present study has several limitations. One limitation of the co-IP assay is that the antibody used for the co-IP experiments does not recognize endogenous mouse LaA to the same extent as the exogenous human LaA, and for this reason, data from wild-type cells are not included in Figure 4D-E. The *Lmna*^−/−^ cells used in this study are known to express a residual truncated form of LaA (Jahn et al., 2012; Kim et al., 2023); however, our current and previous data (Earle et al., 2020) demonstrate full rescue by human LaA, indicating that the fragment does not act in a dominant-negative manner. Additionally, we used LaA antibodies that do not recognize this fragment to avoid confounding effects when assessing exogenous LaA levels and distribution. Finally, we tested only two specific, commonly used N-terminal tags (i.e., FLAG and GFP) on LaA rescue ability, and did not test the effect of inserting different linkers between the tags and LaA. This leaves open the possibility that use of linker residues that confer flexibility, or the use of other N-terminal tags, or internal tags might lead to less severe or no effects on lamin function.

In summary, our data indicate that even when some assays suggest normal function of the tagged protein, this does not exclude that other functions are affected by the addition of a tag, even for small tags. Therefore, non-tagged constructs should be included routinely as controls, and findings using tagged proteins should be confirmed using non-tagged versions of the protein whenever possible.

## Materials and Methods

### Cell culture

Spontaneously immortalized wild-type and *Lmna^−/−^*MEFs were a kind gift from Colin Stewart (Sullivan et al., 1999) and were maintained in DMEM supplemented with 10% FBS and 1% penicillin/streptomycin. Cells were passaged at 80-90% confluency and routinely checked for mycoplasma contamination. For stable genetic manipulations, pseudovirus particles or the Piggybac transposase system were used as described previously (Denais et al., 2016; Earle et al., 2020). Antibiotic selection was performed using Puromycin at 3 µg/ml and Blasticidin at 4.5 µg/ml for at least 1 week. Clonal isolation was performed via serial dilution in a 96-well plate, followed by screening of putative clones by immunofluorescence for homogenous levels of protein expression. Doxycycline titrations were performed in 24-well plates using 1:2 serial dilutions.

### Genetic construct information

The doxycycline-inducible FLAG-hLaA construct has been described previously (Odell et al., 2024). Untagged LaA was cloned via Gibson assembly following digestion of the pPB-rtTA-hCas9-puro-PB backbone (Wang et al., 2017) with Nhe1 and Age1. Untagged human LaA was amplified from pCDH-CMV-hLamin_A-IRES-copGFP-EF1-puro (Earle et al., 2020) using the PCR primers 5’- ACCCTCGTAAAGGTCTAGAGACCATGGAGACCCCGTCC-3’ and 5’- CCGTTTAAACTCATTACTAATTACATGATGCTGCAGTTCTGG-3’. Enhanced GFP-LaA was cloned into the same backbone following PCR amplification using the PCR primers 5’- ACCCTCGTAAAGGTCTAGAGACCATGGTGAGCAAGGGC-3’ and 5’- CCGTTTAAACTCATTACTAATTACATGATGCTGCAGTTCTGG-3’. Following cloning, all constructs were verified with Sanger sequencing of the inserts. The cGAS-mCherry reporter used in this study has been described previously (Denais et al., 2016). For Piggybac transposition, plasmids containing the desired insert were co-transfected with a plasmid encoding a hyperactive transposase (2:1 vector plasmid: hyperactive transposase plasmid) using the Purefection system according to the manufacturer’s instructions.

### Differential protein extraction

Cells (1×10^5^) were seeded in wells of a 6-well plate, and doxycycline was added at the appropriate concentration for 24 h to induce protein expression. To isolate the easily soluble fraction, cells were lysed using 200 µl low-salt buffer (0.5 × PBS, 50 mM HEPES (pH 8.0), 10 mM MgCl2, 1 mM EGTA, 0.2 % NP-40 Alternative). Cells were lysed on ice for 5 minutes, then cells were scraped off the plate, transferred to 1.7 ml microcentrifuge tubes, and spun at 4°C for 5 min at max speed in a benchtop centrifuge. The supernatant was saved as the “soluble fraction”. The pellet was resuspended in 200 µl high-salt RIPA buffer (12 mM Sodium Deoxycholate, 50 mM Tris pH 8.0, 750 mM NaCl, 1% (v/v) NP-40 Alternative, 0.1% (v/v) sodium dodecyl sulfate), vortexed for 5 minutes, sonicated (Branson 450 Digital Sonifier) for 30 s at 36% amplitude, boiled for 2 min, and centrifuged at 4°C for 10 min. The supernatant from this step was saved as the “insoluble fraction”. Equal amounts of each fraction (20 µl) were mixed with 5 × Laemmli Buffer, boiled for 3 min, and then separated by SDS-PAGE as described below.

### Immunofluorescence

Cells were seeded on fibronectin-coated glass coverslips overnight, then doxycycline was added at the appropriate concentration for 24 h to induce protein expression. Fixation was performed with 4% paraformaldehyde in PBS for 15 minutes at room temperature, followed by 3, 5-minute washes with IF wash buffer containing 0.2% Triton X, 0.25% Tween 20, and 0.3% BSA in PBS. Cells were blocked in 3% BSA in PBS for 1 hour, then primary antibodies were added for 1 h in blocking buffer at room temperature or overnight at 4°C. Primary antibodies used: anti-LaA (Millipore MAB3540, 1:250), Emerin (Leica Emerin-NCL, 1:1000). DAPI was added 1:1000 in PBS for 15 minutes. Secondary antibodies used were Alexa Fluor 488 or 568- conjugated donkey anti mouse/rabbit antibodies (Invitrogen) diluted 1:250 in 3% BSA in PBS. Coverslips were mounted on glass slides using mowiol and kept in the dark until imaging.

### Immunoblotting

Cells (1 × 10^5^) were seeded in wells of six-well plates overnight, and then doxycycline was added at the appropriate concentration for 24 h to induce protein expression. Cells were lysed using a high salt RIPA buffer (12 mM Sodium Deoxycholate, 50 mM Tris pH 8.0, 750 mM NaCl, 1% (v/v) NP-40 Alternative, 0.1% (v/v) sodium dodecyl sulfate in ultrapure water). To extract LaA, lysates were vortexed for 5 min, sonicated (Branson 450 Digital Sonifier) for 30 s at 36% amplitude, boiled for 2 min, centrifuged at 4°C for 10 min and stored at –70°C. Protein concentration was determined using Bradford assay. Equal protein amounts were denatured in 5 × Laemmli buffer by boiling for 3 min, loaded onto 4–12% Bis-Tris gels (Invitrogen NP0322), run for 1.5 h at 100 V, then transferred for 1 h at 16 V onto PVDF membrane. Membranes were blocked for 1 h in blocking buffer containing 3% BSA in Tris-buffered saline + 1% Tween 20. Primary antibodies used for Figure 1D: rabbit anti Lamin A/C (Cell Signaling 2032S, 1:1000), mouse anti PCNA (Santa Cruz sc-56, 1:1000). Primary antibodies used for Figure 4D: mouse anti LaA (Millipore MAB3540; 1:3000), mouse anti-Emerin (Leica NCL-Emerin; 1:1000). Secondary antibodies used: Licor IRDye 680RD Donkey anti-Mouse IgG (926-68072) 1:5000, Licor IRDye 800CW Donkey anti-Rabbit IgG (926-32213) 1:5000. Secondary antibodies were added for 1 h at room temperature in blocking buffer, followed by 3 10 min washes. Membranes were imaged using the Odyssey Licor scanner, and then cropped and brightness/ contrast was adjusted using Image Studio Lite (version 5.2) software. Uncropped versions of all blots are available in Suppl. Figure S2.

### Immunoprecipitation

Co-IP studies were performed using the Pierce Protein A/G magnetic beads (ThermoFisher 88802) according to manufacturer’s instructions. Briefly, whole cell lysates were prepared as described above, except the boiling step was omitted to prevent denaturation. Lysates were incubated with anti-LaA antibody (Millipore MAB3540) 1:100 overnight at 4°C with gentle agitation. For control IPs, a non-specific IgG3 (Cell Signaling 37988) was used instead. Magnetic beads (25 µl) were washed 2 times with lysis buffer, then beads were mixed with the lysates and complexes were allowed to form for 1 hour at room temperature with gentle mixing. Bound antibody-antigen complexes were isolated using a magnetic separator, and samples were washed 5 times in IP Wash Buffer containing 50 mM Tris HCl, pH 8.0, 0.3 M NaCl, and 0.3% Triton. Samples were eluted using Laemmli buffer (Biorad 1610737EDU) and boiled for 5 minutes to release bound proteins. The entire volume of eluted protein was loaded onto gels for SDS-PAGE as above, along with 4% lysate input that was set aside prior to addition of primary antibody. To compare the amount of Emerin co-immunoprecipitated with different LaA constructs, we first quantified the intensity of the bands for the co-immunoprecipitated Emerin, Emerin in the IP input, and immunoprecipitated LaA. We then divided the levels of the co-immunoprecipitated Emerin by the levels of Emerin in the input. This fraction was then normalized to the levels of LaA immunoprecipitated in each condition

### Micropipette aspiration assay

Micropipette aspiration was performed according to a previously published protocol (Davidson et al., 2019). In brief, 1-3 × 10^6^ cells were suspended in 2% BSA in PBS supplemented with 0.2% FBS and 10 mM EDTA to prevent cell clumping or adherence. Hoechst 33342 was added 1:1000 immediately before the cell suspension was transferred to the micropipette device. Cells were perfused into the device using the following pressure settings: inlet port (top): 1.0 psi; inlet port (bottom): 0.2 psi. Cells flow through the device and are trapped in the micropipette pockets because of the difference in pressure on the different regions of the device, allowing for aspiration of the nucleus into the small ‘micropipette-like’ aspiration channel. A small pressure gradient drives the perfusion of the cell suspension through the larger channel, with some of the cells becoming deposited in the ‘pockets’ that contain the smaller aspiration channels. Cells are gradually perfused, or at least partially aspirated, into these smaller aspiration channels, driven by a larger pressure gradient across these channels. Once a flow of cells was established in the device, cells were cleared from the pockets to allow new cells to enter, and images were acquired every 5 s for 40 frames. Nuclear protrusion length was measured using a MatLab script available at (https://github.com/Lammerding/MATLAB-micropipette_analysis).

### Microscopy

Confocal images were acquired on a Zeiss LSM900 series confocal microscope with airyscan module using a 40× water immersion objective. The optimal z-slice size was automatically determined using Zen Blue (Zeiss) software. Airy units for images were set between 1.5 and 2.5. Micropipette aspiration data was acquired using an inverted Zeiss Observer Z1 epifluorescence microscope with Hamamatsu Orca Flash 4.0 camera. The image acquisition for micropipette aspiration experiments was automated with Zen Blue (Zeiss) software.

### Image analysis

Nuclear circularity calculations were performed using a FIJI macro described in (Odell et al., 2024). Briefly, this macro performs a background subtraction and thresholds the image based on the DAPI channel to identify nuclei, and then measures the circularity of each nucleus and mean intensity in each channel using the Analyze Particles function.

Intensity profile measurements were performed using a FIJI macro available on request. Briefly, this macro used the “Plot Profile” feature in FIJI to measure the LaA intensity across a line drawn across a z-slice through the center of the nucleus. To account for differences in nuclear size, the intensity profiles are converted into relative nuclear distances, as depicted in Figure 2B.

For measurements of nuclear/cytoplasmic ratios of Emerin, nuclear Emerin was measured using the same macro as the nuclear circularity calculations, and for each cell, cytoplasmic Emerin was obtained by manually drawing a ROI adjacent to the corresponding nucleus for each cell. Then, mean nuclear Emerin intensity was divided by mean cytoplasmic Emerin intensity for each cell.

NE rupture rates were determined based on the fraction of cells with nuclei positive for the NE reporter cGAS-mCherry, as described previously (Denais et al., 2016; Odell et al., 2024). Note that the cGAS-mCherry reporter contains two mutations (E225A/D227A) introduced into the cGAS catalytic domain that prevent interferon production and downstream immune signaling, while still allowing for the protein to bind DNA and localize at NE rupture sites (Denais et al., 2016). Briefly, cells were seeded on fibronectin-coated glass coverslips in wells of a 24-well plate, allowed to adhere overnight, and then LaA expression was induced for 24 h using doxycycline concentrations identified in Figure 1. Cells were fixed as above and DNA was labelled using 4’,6- diamidino-2-phenylindole (DAPI) diluted 1:1000 in PBS. For counts of cGAS-mCherry positive nuclei, images were blinded and nuclei were scored as “cGAS positive” or “cGAS negative” based on the presence or absence of mCherry puncta adjacent to each DAPI stained nucleus. For all image analysis, cells on the edges of the image, dead cells, or mitotic cells were excluded manually from the analysis.

### Statistical analysis and figure generation

All analyses were performed using GraphPad Prism. For comparisons of more than 3 groups, one-way ANOVA with Tukey’s multiple comparison test was performed on the replicate means. For all quantification of cell-level measurements (i.e., Figures 1C, 2B, 2D, 3D, and 4C) grey points represent measurements from individual cells, black points represent replicate means, and bars indicate overall means. Sets of points with the same letter above them are not statistically significantly different, whereas different letters indicate *p* < 0.05. To compare proportions of nuclei with cGAS-mCherry foci, Fisher’s exact test was performed on the total proportion of cGAS-mCherry positive versus negative nuclei. In Figure 3E, individual data points indicate means from independent experiments; bars indicate overall proportions based on three independent experiments. At least 180 nuclei in total were scored for each cell line. Different letters above each set of points indicate *p* < 0.05 based on Fisher’s exact test. The full results (P-values) of the statistical tests are shown in Supplemental Tables 1-8.

Experiments were performed a minimum of three independent times, and for qualitative image analysis, observers were blinded to genotype or treatment conditions when scoring phenotypes. Our statistical analysis was developed in close consultation with the Cornell Statistical Consulting Unit. Figures were assembled using Adobe Illustrator.

## Acknowledgements

We thank the Biotechnology Resource Center (BRC) Flow Cytometry Facility (RRID: SCR_021740) and sequencing facility (RRID: SCR_021727) at the Cornell Institute of Biotechnology for their resources and technical assistance. This work was performed in part at the Cornell NanoScale Science & Technology Facility, a member of the National Nanotechnology Coordinated Infrastructure, which is supported by the National Science Foundation (award NNCI-2025233). This work was supported by awards from the Volkswagen Foundation (A130142 to J.L.), the National Institutes of Health (R01 HL082792, R01 GM137605, and R35 GM153257 to J.L.), and the National Science Foundation (URoL2022048 to J.L.). The content of this manuscript is solely the responsibility of the authors and does not necessarily represent the official views of the National Institutes of Health.

