## Supplementary Materials for "N-terminal tags impair the ability of Lamin A to provide structural support to the nucleus"

### Supplementary Data

#### Supplemental Figures:

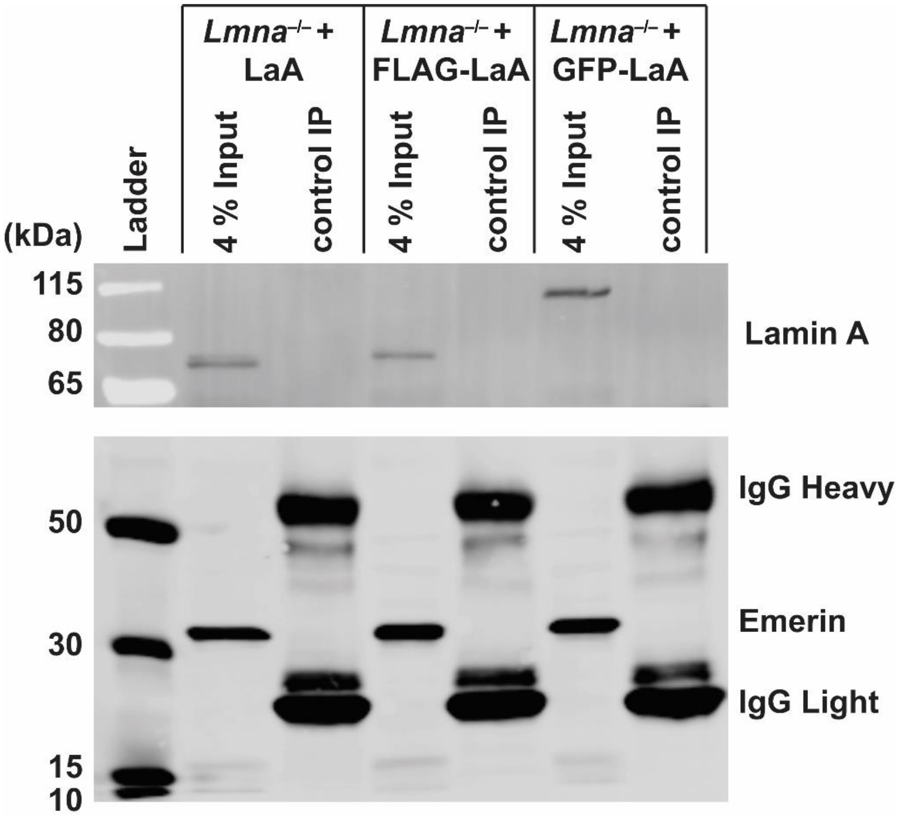


**Supplemental Figure 1: Nonspecific IgG does not immunoprecipitate LaA nor Emerin**. We performed immunoprecipitation experiment analogous to those described in Figure 4D-E, but using a non-specific isotype matched IgG (control IP). Unlike the results obtained with a LaA specific antibody shown in Figure 4D-E, none of the LaA constructs, nor Emerin, were IP'ed using the control IgG, despite being present in the input

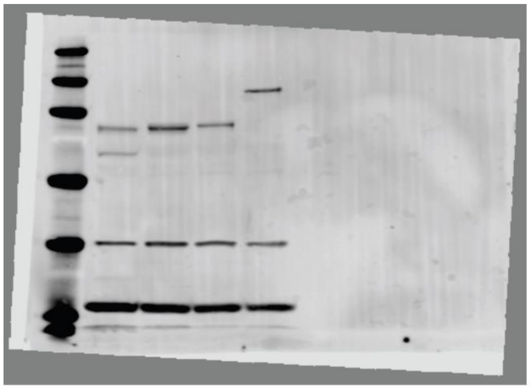

**Full blot for Figure 1D**
**Antibodies used:**
**rabbit anti Lamin A/C (Cell Signaling 2032S)**
**mouse anti PCNA (Santa Cruz sc-56)**
**rabbit anti H3 (Cell Signalling 4499S)**

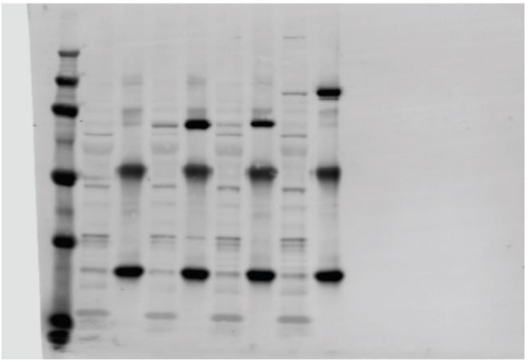

**Full blot for Figure 4D:**
**Antibodies used:**
**mouse anti Lamin A/C (Millipore MAB3540)**
**mouse anti-Emerin (Leica NCL-Emerin)**

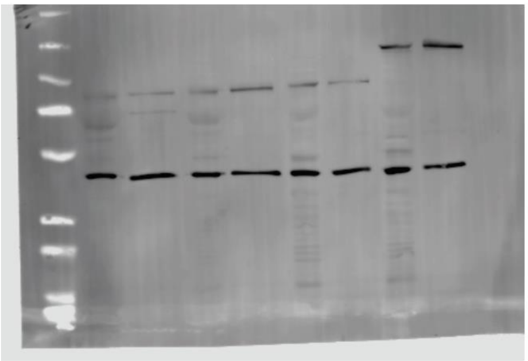

**Full blot for Figure 2E (rabbit)**
**Antibodies used:**
**rabbit anti Lamin A/C (Cell Signaling 2032S)**
**rabbit anti actin (Cell Signalling 8456S)**

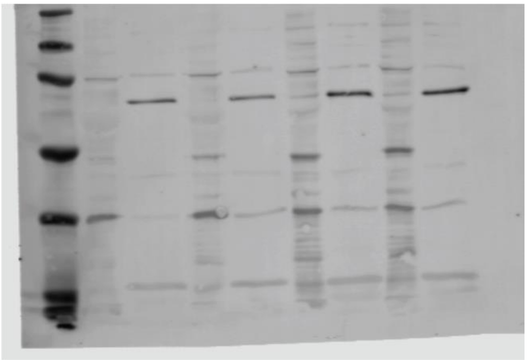

**Full blot for Figure 2E (mouse)**
**Antibodies used:**
**mouse anti Lamin B1 (Santa Cruz sc-374015)**
**mouse anti PCNA (Santa Cruz sc-56)**

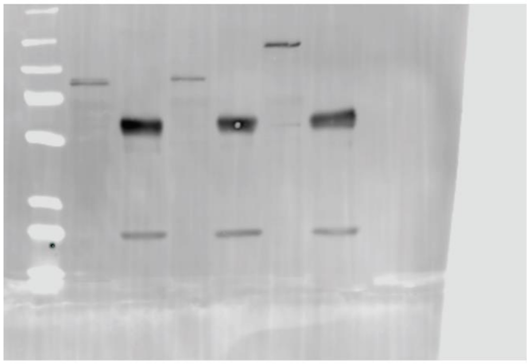

**Full blot for Suppl. Figure 1 (rabbit)**
**Antibodies used:**
**rabbit anti Lamin A/C (Cell Signaling 2032S)**

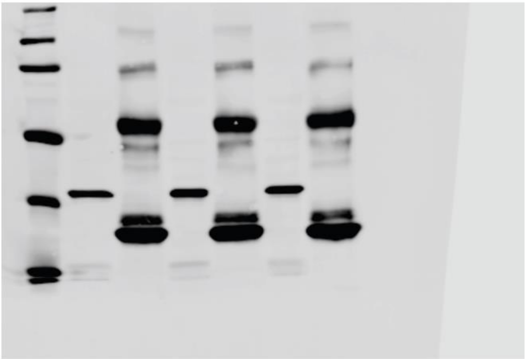

**Full blot for Suppl. Figure 1 (mouse)**
**Antibodies used:**
**mouse anti-Emerin (Leica NCL-Emerin)**

**Supplemental Figure 2: Blot transparency.** Full, uncropped blots are shown for each of the immunoblots presented here. ThermoFisher PageRuler plus prestained protein ladder (26619) was used as molecular weight marker and is present in the first, leftmost lane in all blots.

**Supplemental Tables:**

| **Statistics for Figure 1C:** | | |
|---|---|---|
| **Tukey's multiple comparisons test** | **Adjusted P Value** | **Below threshold?** |
| [LaA no dox] vs. [FLAG-LaA no dox] | >0.9999 | No |
| [LaA no dox] vs. [GFP-LaA no dox] | >0.9999 | No |
| [LaA no dox] vs. [LaA + Dox] | <0.0001 | Yes |
| [LaA no dox] vs. [FLAG-LaA + Dox] | <0.0001 | Yes |
| [LaA no dox] vs. [GFP-LaA + Dox] | <0.0001 | Yes |
| [FLAG-LaA no dox] vs. [GFP-LaA no dox] | >0.9999 | No |
| [FLAG-LaA no dox] vs. [LaA + Dox] | <0.0001 | Yes |
| [FLAG-LaA no dox] vs. [FLAG-LaA + Dox] | <0.0001 | Yes |
| [FLAG-LaA no dox] vs. [GFP-LaA + Dox] | <0.0001 | Yes |
| [GFP-LaA no dox] vs. [LaA + Dox] | <0.0001 | Yes |
| [GFP-LaA no dox] vs. [FLAG-LaA + Dox] | <0.0001 | Yes |
| [GFP-LaA no dox] vs. [GFP-LaA + Dox] | <0.0001 | Yes |
| [LaA + Dox] vs. [FLAG-LaA + Dox] | >0.9999 | No |
| [LaA + Dox] vs. [GFP-LaA + Dox] | 0.9967 | No |
| [FLAG-LaA + Dox] vs. [GFP-LaA + Dox] | 0.994 | No |

**Supplemental Table 1**: Results of Tukey's multiple comparisons test performed on data presented in Figure 1C

| **Statistics for Figure 2B:** | | |
|---|---|---|
| **Tukey's multiple comparisons test** | **Adjusted P Value** | **Below threshold?** |
| [LaA no dox] vs. [FLAG-LaA no dox] | 0.4548 | No |
| [LaA no dox] vs. [GFP-LaA no dox] | 0.4279 | No |
| [LaA no dox] vs. [LaA + Dox] | <0.0001 | Yes |
| [LaA no dox] vs. [FLAG-LaA + Dox] | <0.0001 | Yes |
| [LaA no dox] vs. [GFP-LaA + Dox] | <0.0001 | Yes |
| [LaA no dox] vs. [WT] | <0.0001 | Yes |
| [FLAG-LaA no dox] vs. [GFP-LaA no dox] | >0.9999 | No |
| [FLAG-LaA no dox] vs. [LaA + Dox] | 0.0003 | Yes |
| [FLAG-LaA no dox] vs. [FLAG-LaA + Dox] | 0.0003 | Yes |
| [FLAG-LaA no dox] vs. [GFP-LaA + Dox] | 0.0003 | Yes |
| [FLAG-LaA no dox] vs. [WT] | <0.0001 | Yes |
| [GFP-LaA no dox] vs. [LaA + Dox] | 0.0003 | Yes |
| [GFP-LaA no dox] vs. [FLAG-LaA + Dox] | 0.0003 | Yes |
| [GFP-LaA no dox] vs. [GFP-LaA + Dox] | 0.0003 | Yes |
| [GFP-LaA no dox] vs. [WT] | <0.0001 | Yes |
| [LaA + Dox] vs. [FLAG-LaA + Dox] | >0.9999 | No |
| [LaA + Dox] vs. [GFP-LaA + Dox] | >0.9999 | No |

| | | |
|---|---|---|
| [LaA + Dox] vs. [WT] | 0.1011 | No |
| [FLAG-LaA + Dox] vs. [GFP-LaA + Dox] | >0.9999 | No |
| [FLAG-LaA + Dox] vs. [WT] | 0.0829 | No |
| [GFP-LaA + Dox] vs. [WT] | 0.0883 | No |

**Supplemental Table 2:** Results of Tukey's multiple comparisons test performed on data presented in Figure 2B

| **Statistics for Figure 2D:** | | |
|---|---|---|
| **Tukey's multiple comparisons test** | **Adjusted P Value** | **Below threshold?** |
| [LaA + Dox] vs. [FLAG-LaA + Dox] | 0.0016 | Yes |
| [LaA + Dox] vs. [GFP-LaA + Dox] | 0.0017 | Yes |
| [LaA + Dox] vs. [WT] | 0.8025 | No |
| [FLAG-LaA + Dox] vs. [GFP-LaA + Dox] | >0.9999 | No |
| [FLAG-LaA + Dox] vs. [WT] | 0.0047 | Yes |
| [GFP-LaA + Dox] vs. [WT] | 0.0048 | Yes |

**Supplemental Table 3:** Results of Tukey's multiple comparisons test performed on data presented in Figure 2D

| **Statistics for Figure 2F:** | | |
|---|---|---|
| **Tukey's multiple comparisons test** | **Adjusted P Value** | **Below threshold?** |
| [LaA + Dox] vs. [WT] | 0.7781 | No |
| [FLAG-LaA + Dox] vs. [WT] | <0.0001 | Yes |
| [GFP-LaA + Dox] vs. [WT] | 0.0008 | Yes |
| [LaA + Dox] vs. [FLAG-LaA + Dox] | <0.0001 | Yes |
| [LaA + Dox] vs. [GFP-LaA + Dox] | 0.0021 | Yes |
| [FLAG-LaA + Dox] vs. [GFP-LaA + Dox] | <0.0001 | Yes |

**Supplemental Table 4:** Results of Tukey's multiple comparisons test performed on data presented in Figure 2F

| **Statistics for Figure 3D:** | | |
|---|---|---|
| **Tukey's multiple comparisons test** | **Adjusted P Value** | **Below threshold?** |
| [LaA no dox] vs. [WT] | <0.0001 | Yes |
| [FLAG-LaA no dox] vs. [WT] | <0.0001 | Yes |
| [GFP-LaA no dox] vs. [WT] | <0.0001 | Yes |
| [LaA + Dox] vs. [WT] | 0.6947 | No |
| [FLAG-LaA + Dox] vs. [WT] | 0.0012 | Yes |
| [GFP-LaA + Dox] vs. [WT] | 0.0002 | Yes |
| [LaA no dox] vs. [FLAG-LaA no dox] | 0.9653 | No |
| [LaA no dox] vs. [GFP-LaA no dox] | 0.9897 | No |
| [LaA no dox] vs. [LaA + Dox] | <0.0001 | Yes |
| [LaA no dox] vs. [FLAG-LaA + Dox] | 0.0003 | Yes |

| | | |
|---|---|---|
| [LaA no dox] vs. [GFP-LaA + Dox] | 0.0022 | Yes |
| [FLAG-LaA no dox] vs. [GFP-LaA no dox] | >0.9999 | No |
| [FLAG-LaA no dox] vs. [LaA + Dox] | <0.0001 | Yes |
| [FLAG-LaA no dox] vs. [FLAG-LaA + Dox] | <0.0001 | Yes |
| [FLAG-LaA no dox] vs. [GFP-LaA + Dox] | 0.0005 | Yes |
| [GFP-LaA no dox] vs. [LaA + Dox] | <0.0001 | Yes |
| [GFP-LaA no dox] vs. [FLAG-LaA + Dox] | 0.0001 | Yes |
| [GFP-LaA no dox] vs. [GFP-LaA + Dox] | 0.0006 | Yes |
| [LaA + Dox] vs. [FLAG-LaA + Dox] | 0.0208 | Yes |
| [LaA + Dox] vs. [GFP-LaA + Dox] | 0.0027 | Yes |
| [FLAG-LaA + Dox] vs. [GFP-LaA + Dox] | 0.9102 | No |

**Supplemental Table 5:** Results of Tukey’s multiple comparisons test performed on data presented in Figure 3D

| **Statistics for Figure 3F:** | | |
|---|---|---|
| **Fisher's exact test:** | **P value** | **Below threshold?** |
| [LaA no dox] vs. [WT] | <0.0001 | Yes |
| [FLAG-LaA no dox] vs. [WT] | <0.0001 | Yes |
| [GFP-LaA no dox] vs. [WT] | <0.0001 | Yes |
| [LaA + Dox] vs. [WT] | 0.069 | No |
| [FLAG-LaA + Dox] vs. [WT] | 0.3357 | No |
| [GFP-LaA + Dox] vs. [WT] | 0.2858 | No |
| [LaA no dox] vs. [FLAG-LaA no dox] | 0.6299 | No |
| [LaA no dox] vs. [GFP-LaA no dox] | 0.5019 | No |
| [LaA no dox] vs. [LaA + Dox] | 0.0187 | Yes |
| [LaA no dox] vs. [FLAG-LaA + Dox] | 0.001 | Yes |
| [LaA no dox] vs. [GFP-LaA + Dox] | 0.0041 | Yes |
| [FLAG-LaA no dox] vs. [GFP-LaA no dox] | 0.8733 | No |
| [FLAG-LaA no dox] vs. [LaA + Dox] | 0.0434 | Yes |
| [FLAG-LaA no dox] vs. [FLAG-LaA + Dox] | 0.0017 | Yes |
| [FLAG-LaA no dox] vs. [GFP-LaA + Dox] | 0.0081 | Yes |
| [GFP-LaA no dox] vs. [LaA + Dox] | 0.025 | Yes |
| [GFP-LaA no dox] vs. [FLAG-LaA + Dox] | 0.002 | Yes |
| [GFP-LaA no dox] vs. [GFP-LaA + Dox] | 0.0171 | Yes |
| [LaA + Dox] vs. [FLAG-LaA + Dox] | 0.4572 | No |
| [LaA + Dox] vs. [GFP-LaA + Dox] | 0.6158 | No |
| [FLAG-LaA + Dox] vs. [GFP-LaA + Dox] | >0.9999 | No |

**Supplemental Table 6:** Results of Fisher’s exact test performed on data presented in Figure 3F

| **Statistics for Figure 4C:** |
|---|

| Tukey's multiple comparisons test | Adjusted P Value | Below threshold? |
|---|---|---|
| [LaA no dox] vs. [LaA + Dox] | 0.0001 | Yes |
| [LaA no dox] vs. [FLAG-LaA no dox] | 0.9887 | No |
| [LaA no dox] vs. [FLAG-LaA + Dox] | 0.0003 | Yes |
| [LaA no dox] vs. [GFP-LaA no dox] | >0.9999 | No |
| [LaA no dox] vs. [GFP-LaA + Dox] | 0.0129 | Yes |
| [LaA no dox] vs. [WT] | <0.0001 | Yes |
| [FLAG-LaA no dox] vs. [LaA + Dox] | 0.0004 | Yes |
| [LaA + Dox] vs. [FLAG-LaA + Dox] | 0.9964 | No |
| [GFP-LaA no dox] vs. [LaA + Dox] | 0.0001 | Yes |
| [LaA + Dox] vs. [GFP-LaA + Dox] | 0.0167 | Yes |
| [LaA + Dox] vs. [WT] | 0.3198 | No |
| [FLAG-LaA no dox] vs. [FLAG-LaA + Dox] | 0.001 | Yes |
| [FLAG-LaA no dox] vs. [GFP-LaA no dox] | 0.9889 | No |
| [FLAG-LaA no dox] vs. [GFP-LaA + Dox] | 0.0381 | Yes |
| [FLAG-LaA no dox] vs. [WT] | <0.0001 | Yes |
| [GFP-LaA no dox] vs. [FLAG-LaA + Dox] | 0.0003 | Yes |
| [FLAG-LaA + Dox] vs. [GFP-LaA + Dox] | 0.0479 | Yes |
| [FLAG-LaA + Dox] vs. [WT] | 0.1312 | No |
| [GFP-LaA no dox] vs. [GFP-LaA + Dox] | 0.0129 | Yes |
| [GFP-LaA no dox] vs. [WT] | <0.0001 | Yes |
| [GFP-LaA + Dox] vs. [WT] | 0.0003 | Yes |

**Supplemental Table 7:** Results of Tukey's multiple comparisons test performed on data presented in Figure 4C

| Statistics for Figure 4E: | | |
|---|---|---|
| **Tukey's multiple comparisons test** | **Adjusted P Value** | **Below threshold?** |
| [LaA + Dox] vs. [FLAG-LaA + Dox] | 0.0451 | Yes |
| [LaA + Dox] vs. [GFP-LaA + Dox] | 0.0314 | Yes |
| [FLAG-LaA + Dox] vs. [GFP-LaA + Dox] | 0.952 | No |

**Supplemental Table 8:** Results of Tukey's multiple comparisons test performed on data presented in Figure 4E.
